# Pseudorabies Virus Infection Accelerates Degradation of the Kinesin-3 Motor KIF1A

**DOI:** 10.1101/845289

**Authors:** Hao Huang, Orkide O. Koyuncu, Lynn W. Enquist

**Affiliations:** Department of Molecular Biology, Princeton University, Princeton, New Jersey, USA

**Keywords:** alphaherpesvirus, pseudorabies virus, kinesin, KIF1A, proteasomal degradation, axonal sorting.

## Abstract

Alphaherpesviruses, including pseudorabies virus (PRV), are neuroinvasive pathogens that establish life-long latency in peripheral ganglia following the initial infection at mucosal surfaces. The establishment of latent infection and the subsequent reactivations during which newly-assembled virions are sorted into and transported anterogradely inside axons to the initial mucosal site of infection, rely on axonal bidirectional transport mediated by microtubule-based motors. Previous studies using cultured peripheral nervous system (PNS) neurons have demonstrated that KIF1A, a kinesin-3 motor, mediates the efficient axonal sorting and transport of newly-assembled PRV virions. In this study, we report that KIF1A, unlike other axonal kinesins, is an intrinsically unstable protein prone to proteasomal degradation. Interestingly, PRV infection of neuronal cells leads not only to a non-specific depletion of KIF1A mRNA, but also to an accelerated proteasomal degradation of KIF1A proteins, leading to a near depletion of KIF1A protein late in infection. Using a series of PRV mutants deficient in axonal sorting and anterograde spread, we identified the PRV US9/gE/gI protein complex as a viral factor facilitating the proteasomal degradation of KIF1A proteins. Moreover, by using compartmented neuronal cultures that fluidically and physically separate axons from cell bodies, we found that the proteasomal degradation of KIF1A occurs in axons during infection. We propose that PRV anterograde sorting complex, gE/gI/US9, recruits KIF1A to viral transport vesicles for axonal sorting and transport, and eventually accelerates the proteasomal degradation of KIF1A in axons.

**Importance:** Pseudorabies virus (PRV) is an alphaherpesvirus related to human pathogens herpes simplex virus −1, −2 and varicella zoster virus. Alphaherpesviruses are neuroinvasive pathogens that establish life-long latent infections in the host peripheral nervous system (PNS). Following reactivation from latency, infection spreads from the PNS back via axons to the peripheral mucosal tissues, a process mediated by kinesin motors. Here, we unveil and characterize the underlying mechanisms for a PRV-induced, accelerated degradation of KIF1A, a kinesin-3 motor promoting the sorting and transport of PRV virions in axons. We show that PRV infection disrupts the synthesis of KIF1A, and simultaneously promotes the degradation of intrinsically unstable KIF1A proteins by proteasomes in axons. Our work implies that the timing of motor reduction after reactivation would be critical because progeny particles would have a limited time window for sorting into and transport in axons for further host-to-host spread.

## Introduction

Alphaherpesviruses, including human pathogens herpes simplex virus-1 and −2, varicella-zoster virus, and the veterinary pathogen pseudorabies virus (PRV), are neuroinvasive pathogens that establish and maintain life-long latency in the sensory and automatic ganglia of the peripheral nervous system (PNS) of their mammalian hosts following initial infection at mucosal surfaces (1–3). Latently infected neurons in peripheral ganglia undergo occasional reactivation, during which viral genomes are actively replicated and new viral particles assembled. Spread of infection to new hosts requires that these progeny virions be transported to the initial site of epithelial infection, where replication and subsequent host-to-host transmission occur. On rare occasions, infection spreads from the PNS to the central nervous system, often with serious consequences such as herpes simplex encephalitis (HSE).

Axonal bidirectional transport mechanisms are crucial for the virions and virus proteins to move efficiently through the PNS axons, which are often centimeters to meters long (4). In uninfected neurons, microtubule-based motors, including minus-ended directed cytoplasmic dynein and plus-ended directed kinesin superfamily motors, mediate fast axonal transport of RNAs, proteins, organelles, and vesicles between cell bodies and axonal termini for the maintenance and proper functioning of axons (5, 6). Neuroinvasive viruses such as the alphaherpesviruses and rabies virus have evolved to adopt existing axonal transport machinery for efficient neuroinvasion and subsequent spread to new hosts. A notable example is rabies virus (RABV), whose phosphoprotein (P) binds to LC8 dynein light chain, and was shown to hijack the P75NTR-dependent transport to facilitate fast retrograde axonal transport in sensory DRG neurons (7).

Initially after infection of peripheral tissues, alphaherpesvirus virions bind to PNS axon terminals, and viral capsids are released into axons where they recruit cytoplasmic dynein, via inner tegument proteins UL36 and UL37 (8, 9). This complex then moves in the MT minus-end direction towards the cell body and deposits the viral DNA inside the nucleus. After reactivation of a latent PNS infection, kinesin-superfamily motors are recruited both for viral egress at cell bodies, as well as for sorting into axons and subsequent transport of viral structural proteins and newly-assembled virions (10–14). Previous studies in sympathetic PNS neurons revealed that a subpopulation of fully-enveloped PRV virions in transport vesicles recruit KIF1A, a kinesin-3 motor, through the interactions with a gE/gI/US9 tripartite membrane protein complex in lipid rafts at late Golgi network compartments (e.g., late endosomes or the trans-Golgi network (TGN)) (14, 15). After this interaction, fully enveloped virions in transport vesicles are sorted into axons followed by unidirectional anterograde transport before egressing along the axonal shaft (16). In PRV infected neurons, ectopic expression of dominant-negative KIF1A proteins significantly reduced the number of PRV particles transported in the anterograde direction, demonstrating the critical role of KIF1A in sorting virus particles into axons and also in subsequent transport (14). In this report, we focus on a curious observation made by Kramer et al, who noted that the amount of KIF1A in infected cultured neuronal cells decreased significantly late after PRV infection (14).

In uninfected neurons, the kinesin motor KIF1A mediates fast axonal transport of membranous vesicles including synaptic vesicle precursors and dense core vesicles, and is critical in dendrite morphogenesis and synaptogenesis (17–19). Neurons lacking KIF1A showed impaired transport in synaptic vesicle precursors, marked neuronal degeneration, and death in both cultured neurons and *in vivo* (20). Despite its functional importance, several lines of evidence suggest that KIF1A motors are degraded upon the completion of their axonal transportation runs, and are not recycled for further use. Studies with a ligature applied to the mouse sciatic nerve revealed increased antibody staining of KIF1A in the proximal (close to the cell body), but not distal (close to the axonal terminal), region of ligature, suggesting that once KIF1A motors entered axons, they did not return to the cell bodies (e.g. with dyneins) (21). UNC-104, the KIF1A homolog in *C. elegans*, was shown to be degraded by ubiquitin-related pathways at synapses of mechanosensory neurons (22). More importantly, the stability of UNC-104 protein strongly correlates with the motor’s specific binding to cargo, implying a potential mechanistic link between motor degradation and cargo release (22).

In this study, we investigated KIF1A protein dynamics during PRV infection both in neuronal cell lines and in primary neurons. Here, we report that KIF1A (kinesin-3), unlike kinesin-1 and kinesin-2 motors, is intrinsically prone to proteasomal degradation in uninfected neuronal cells. We further showed that PRV infection blocks new synthesis of this motor, and that viral anterograde-sorting complex US9/gE/gI specifically targets KIF1A for proteasomal degradation, resulting in a near depletion of the protein in 24 hours following infection. Using compartmented neuronal cultures that fluidically and physically separate axons from cell bodies, we found that the proteasomal degradation of KIF1A proteins occurs in axons. Moreover, using a series of PRV mutants deficient in anterograde spread, and using adenovirus vectors to express the viral proteins in neuronal cells, we discovered that the PRV US9/gE/gI protein complex targets specifically KIF1A motors for proteasomal degradation. Altogether, these results suggest that progeny PRV particles recruit KIF1A motors for efficient axonal sorting and anterograde transport that leads to the degradation of the motor protein in axons. Since new synthesis of KIF1A is also blocked by PRV infection, these results suggest progeny particles have a limited time window for sorting into and transport in axons for further host-to-host spread.

## Results

### In uninfected, differentiated PC12 cells, the steady state concentration of KIF1A protein is achieved through rapid protein synthesis and proteasomal degradation

We tested the hypothesis that in differentiated PC12 cells axonal kinesins undergo rapid degradation while protein synthesis continues. To determine the role of the proteasome in this process, we treated uninfected differentiated PC12 cells with the proteasomal inhibitor MG132 for 2, 4, 6, and 8 hours, and monitored the protein concentration of three different kinesin motors: KIF5 (kinesin-1, and the conventional kinesin), KIF3A (kinesin-2), and KIF1A (kinesin-3). MG132 treatment led to more than a 2-fold increase in KIF1A concentration within 6 hours post treatment, indicating that KIF1A protein rapidly accumulated upon proteasome inhibition (Figure 1A and B). However, this surge in KIF1A concentration was ablated when PC12 cells were simultaneously treated with MG132 and cycloheximide, a protein synthesis inhibitor, and KIF1A concentration remained at the similar levels as in untreated cells. These experiments demonstrated that KIF1A proteins are degraded by the proteasomes with new proteins efficiently made to maintain protein steady state levels. Consistent with this idea, in cells treated with CHX alone, KIF1A concentration dropped by over 50% within 4 hours of treatment (Figure 1A and B).

**Fig. 1.**
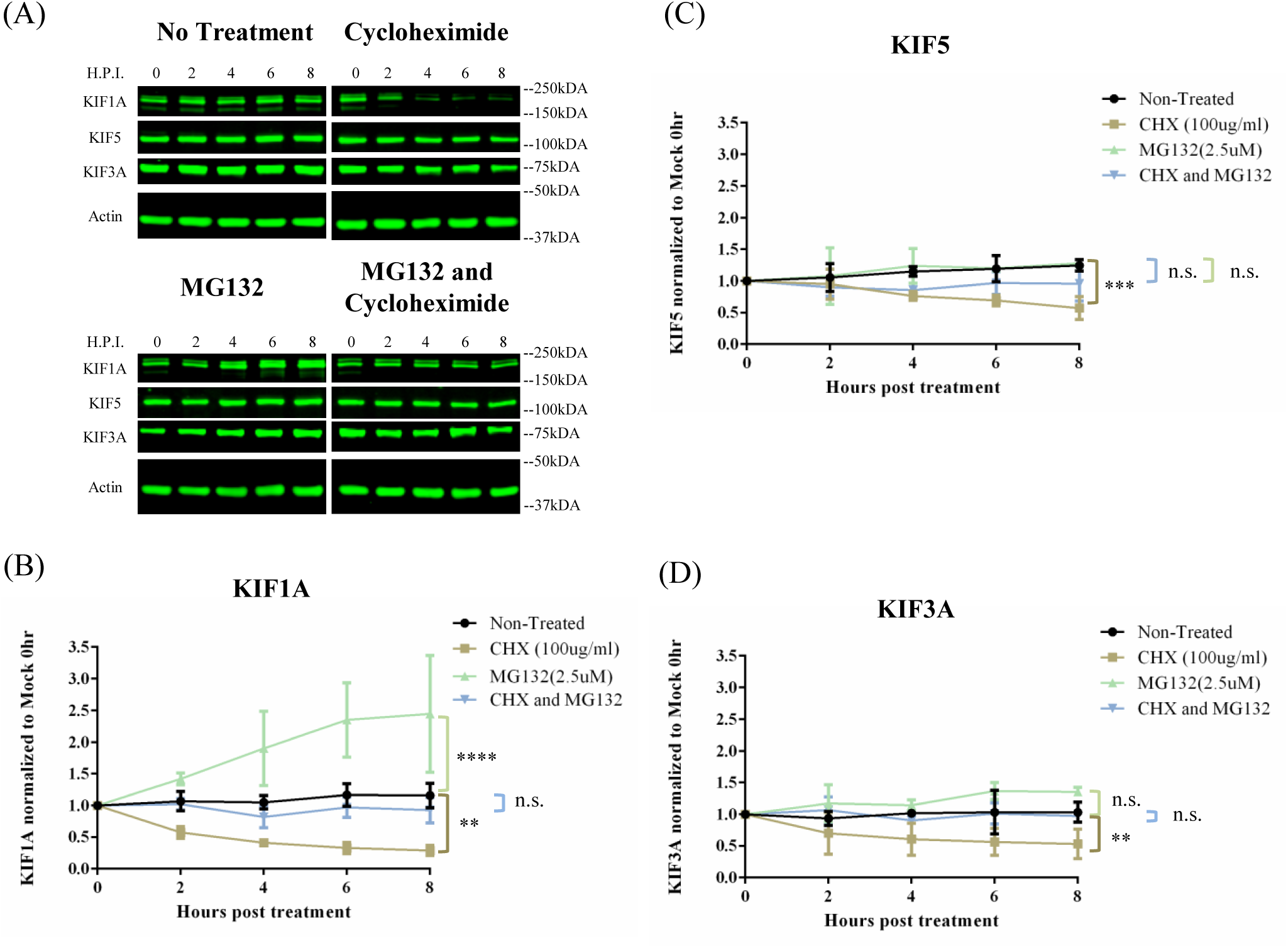
KIF1A protein undergoes rapid accumulation and degradation in neuronal cells upon MG132 and CHX treatments. (A) Differentiated PC12 cells were treated with the translation inhibitor cycloheximide, proteasome inhibitor MG132, or both cycloheximide and MG132 for 2, 4, 6, 8, hours. Cells were harvested at indicated time points and lysates were analyzed by Western blot to monitor protein levels of KIF1A, KIF5, KIF3A, and actin. (B-D) KIF1A, KIF5, and KIF3A protein levels were measured by band intensities and normalized to actin protein levels for each time interval. Normalized values were again normalized to that of Mock 0 hour samples. Values are means plus SEMs (error bar) from three independent experiments. n.s., not statistically significant, **, P<0.01, ***, P<0.001, ****, P<0.0001 for the indicated comparison at 8 hours post treatment.

In contrast to KIF1A, neither KIF5 nor KIF3A protein concentration increased significantly during MG132 treatment (Figure 1A, C and D). With both MG132 and CHX treatments, KIF5 and KIF3A concentrations appeared unchanged in the 8 h treatment period (Figure 1A, C and D), suggesting that while both proteins are degraded by proteasomes, their rate of degradation is much slower than that of KIF1A. As a result, the half-lives of both KIF5 and KIF3A proteins were much longer (> 8 hours) than the half-life of KIF1A in CHX-treated PC12 cells.

### PRV infection induces specific reduction of KIF1A protein late in infection

The inherent instability of KIF1A in uninfected cells prompted us to explore an observation previously made in our laboratory: the concentration of KIF1A protein, the major kinesin facilitating axonal sorting and transport of PRV virions, decreased late in PRV infection of sympathetic neuronal cells. This observation might explain the significant numbers of immobile PRV particles in axons late in infection (23). Accordingly, we infected differentiated PC12 cells with PRV Becker at an MOI of 20 for 3, 8, 12, 16, 20, and 24 hours, and monitored KIF1A protein levels by quantitative western blot methods. Similar to the previous report, we found that the KIF1A protein concentration dropped by ∼90% at 24 hpi (Figure 2A and B, Figure S1). To determine if PRV infection affects the steady state concentrations of other kinesin motors, we monitored the protein levels of KIF5 and KIF3A, two kinesins well-characterized in mediating axonal transport. Neither KIF5 nor KIF3A protein levels were affected by PRV infection (Figure 2).

**Fig. 2.**
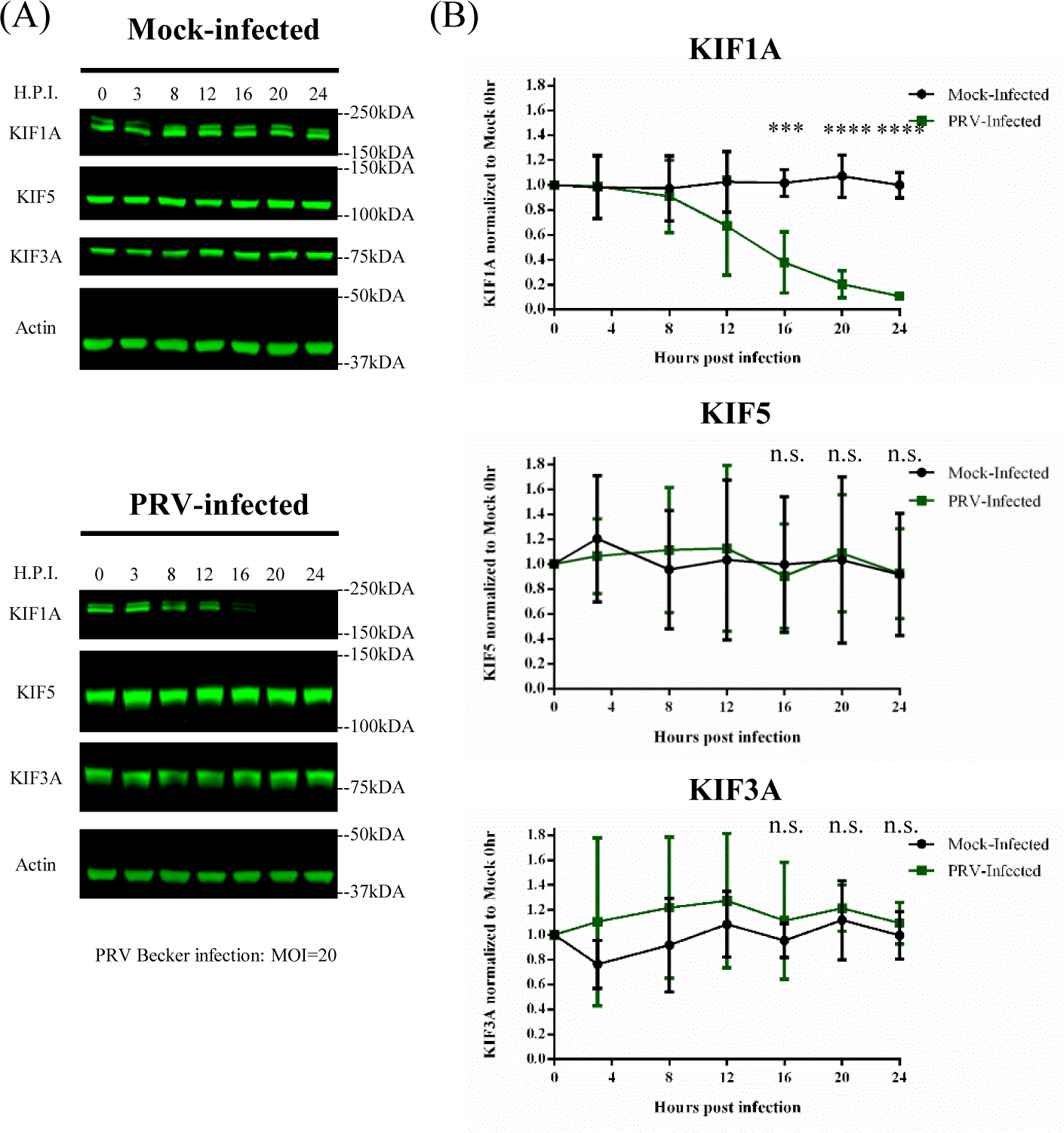
KIF1A protein concentration is reduced during PRV infection in neuronal cells. (A and B) Differentiated PC12 cells were mock-infected or infected with PRV Becker for 3, 8, 12, 16, 20, 24 hours with MOI of 20. Cells were harvested at the time intervals shown, and lysates were analyzed by Western blot to monitor protein levels of KIF1A, KIF5, KIF3A, and actin. Kinesin levels were measured by band intensities and normalized with respect to actin levels at each time interval. Normalized values were again normalized to that of Mock 0 hour samples. Values are means plus SEMs (error bars) from four independent experiments. n.s., not statistically significant, ***, P<0.001, ****, P<0.0001.

### PRV infection reduces both KIF1A and KIF5B transcripts

Alphaherpesviruses promote the instability of many host cell transcripts early in infection, leading to reduced protein synthesis (24–27). We hypothesized that the loss of KIF1A proteins in PRV-infected PC12 cells resulted from such mRNA degradation. We therefore measured the amounts of KIF1A transcripts in PRV infected cells by qRT-PCR. Differentiated PC12 cells were infected with PRV Becker and the attenuated vaccine strain PRV Bartha at an MOI of 20 for 3, 8, 12, 16, 20, 24 hours. Infected cells were collected for RNA extraction and qRT-PCR analysis of KIF1A, KIF5B, and GAPDH mRNA level (Figure 3). In PRV Becker-infected cells, we detected a significant and uniform reduction of KIF1A, KIF5B, and GAPDH transcripts as early as 8 hpi, and their respective concentrations dropped by more than 80% at 16 hpi (Figure 3A).

**Fig. 3.**
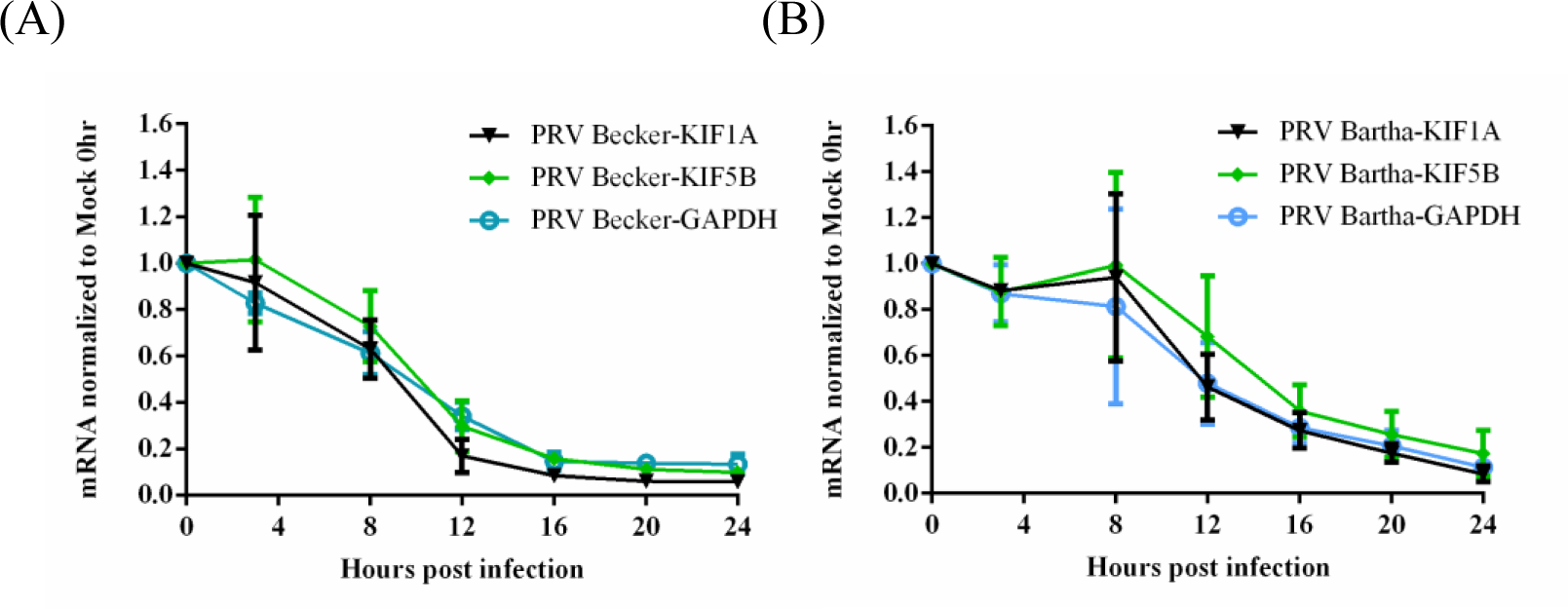
The abundance of KIF1A, KIF5B, and GAPDH transcripts is significantly reduced in PRV-infected neuronal cells. Differentiated PC12 cells were infected with (A) PRV Becker or (B) PRV Bartha for 3, 8, 12, 16, 20, 24 hours with MOI of 20. Cells were harvested at the time intervals shown. Total RNA were extracted and qRT-PCR quantification was performed to measure KIF1A, KIF5B, and GAPDH mRNA levels using 18S rRNA as internal control at each time interval. The normalized values are then normalized to that of Mock 0 hour samples. Values are means plus SEMs (error bars) from three independent experiments.

Compared to PRV Becker infection, PRV Bartha infection decreased KIF1A transcripts at a slower rate, starting at 12 hpi, with more than 80% loss at 20 hpi (Figure 3B). Reduction of GAPDH and KIF5B transcripts occurred with comparable rates and magnitude to the KIF1A transcripts. These experiments showed that KIF1A transcripts were reduced in PRV-infected PC12 cells, but with different kinetics after infection by different PRV strains. Such transcriptional regulation may be one reason why KIF1A protein levels drop significantly during PRV infection. However, this effect was not specific for KIF1A, because KIF5B transcripts were also reduced after infection. Therefore, the specific loss of KIF1A and not KIF5B proteins in infected neurons is not due to reduction of transcript levels.

### KIF1A is degraded by the proteasome during PRV infection in differentiated PC12 cells and in primary SCG neurons

We found that KIF1A (but not other kinesins) is rapidly degraded by the proteasome in uninfected PC12 cells. We also knew from previous studies that complexes of PRV US9 and KIF1A contain the E3-ubiquitin ligases NEDD4 and EP0, suggesting that KIF1A may be degraded via the ubiquitin-proteasome pathway after PRV infection (14). To examine this possibility, we infected differentiated PC12 cells with PRV Becker at an MOI of 20, followed by treatment with the proteasomal inhibitor MG132 at 12 hpi. In mock-infected cells, KIF1A proteins accumulated rapidly, doubling the concentration in 8 hours, after MG132 treatment (Figure 4A and C). However, in PRV Becker infected cells, MG132 treatment led to an effective stabilization of KIF1A protein levels where the protein concentration remained mostly unchanged during the treatment period (Figure 4B and C). This finding suggests that, in the late phase of infection, new KIF1A synthesis is blocked but proteasomal degradation continues, leading to the accelerated loss of KIF1A during PRV infection.

**Fig. 4.**
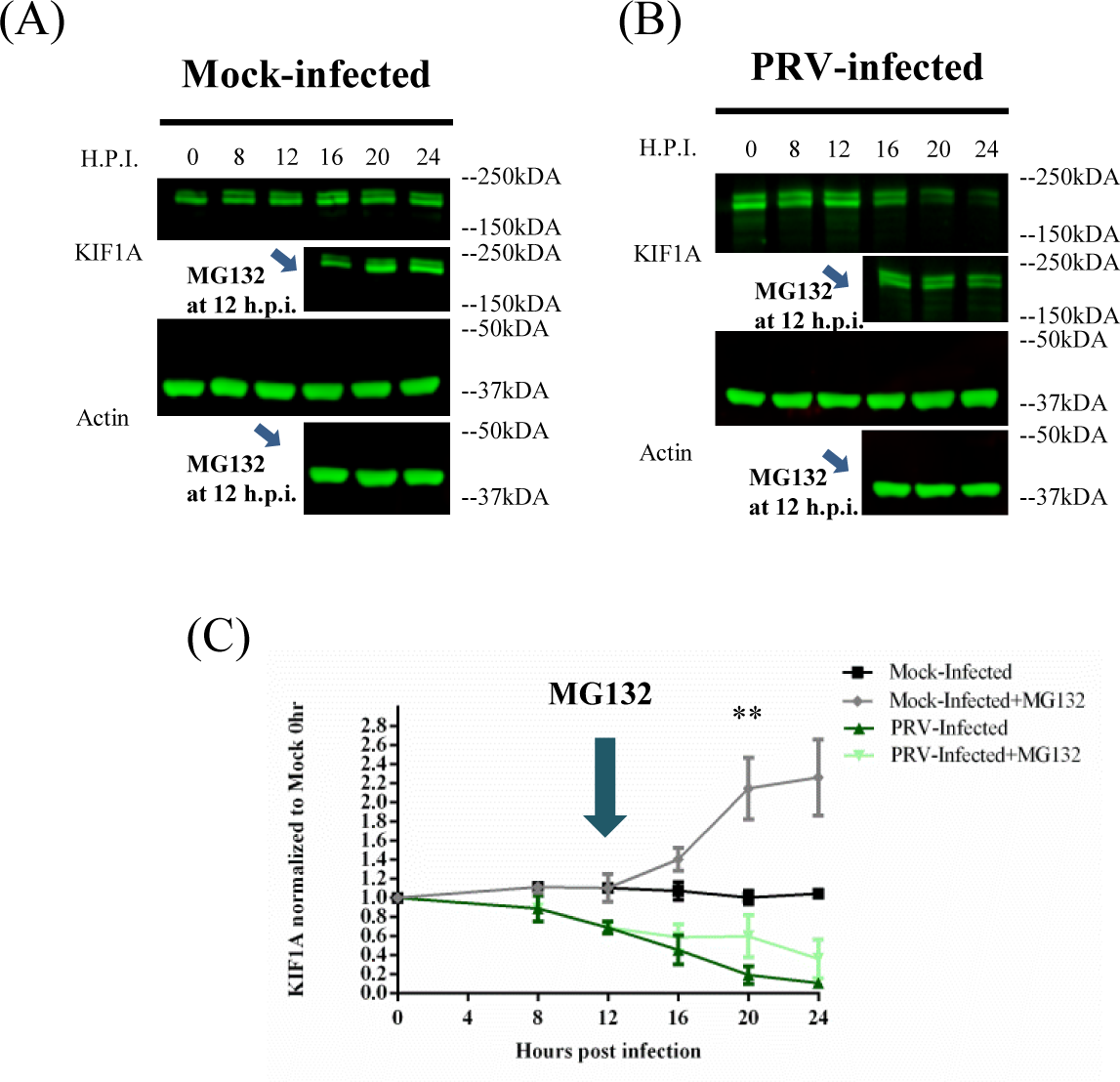
KIF1A is degraded by the proteasome in the late phase of PRV infection. Differentiated PC12 cells were (A) mock-infected or (B) infected with PRV Becker for 8, 12, 16, 20, 24 hours with MOI of 20. In a parallel experiment, the proteasomal inhibitor, MG132, was added at 12 hours post infection at the concentration of 2.5 μM. KIF1A levels were analyzed by Western blot and normalized to actin levels at each time interval. Normalized values were again normalized to that of Mock 0 hour samples. Values are means plus SEMs (error bars) from three independent experiments. **, P<0.01 for comparison between PRV-infected and PRV- infected+MG132 samples.

These studies were done in differentiated PC12 cells that have similar attributes to sympathetic neurons. We performed similar experiments in primary rat superior cervical ganglionic neurons (SCG) cultured in compartmented chambers where we could examine kinesin motor concentration in isolated axons. We cultured primary sympathetic rat SCG neurons in trichambers, in which axons (in the neurite compartment, N) were fluidically and physically separated from cell bodies (in the soma compartment, S) (Figure 5A). SCG cell bodies in the S compartment were infected with PRV Becker for 24 hours and extracts from the S and N compartments were collected for western blot analysis. The concentration of KIF1A protein decreased dramatically both in cell bodies and axons after PRV infection (Figure 5B).

**Fig. 5.**
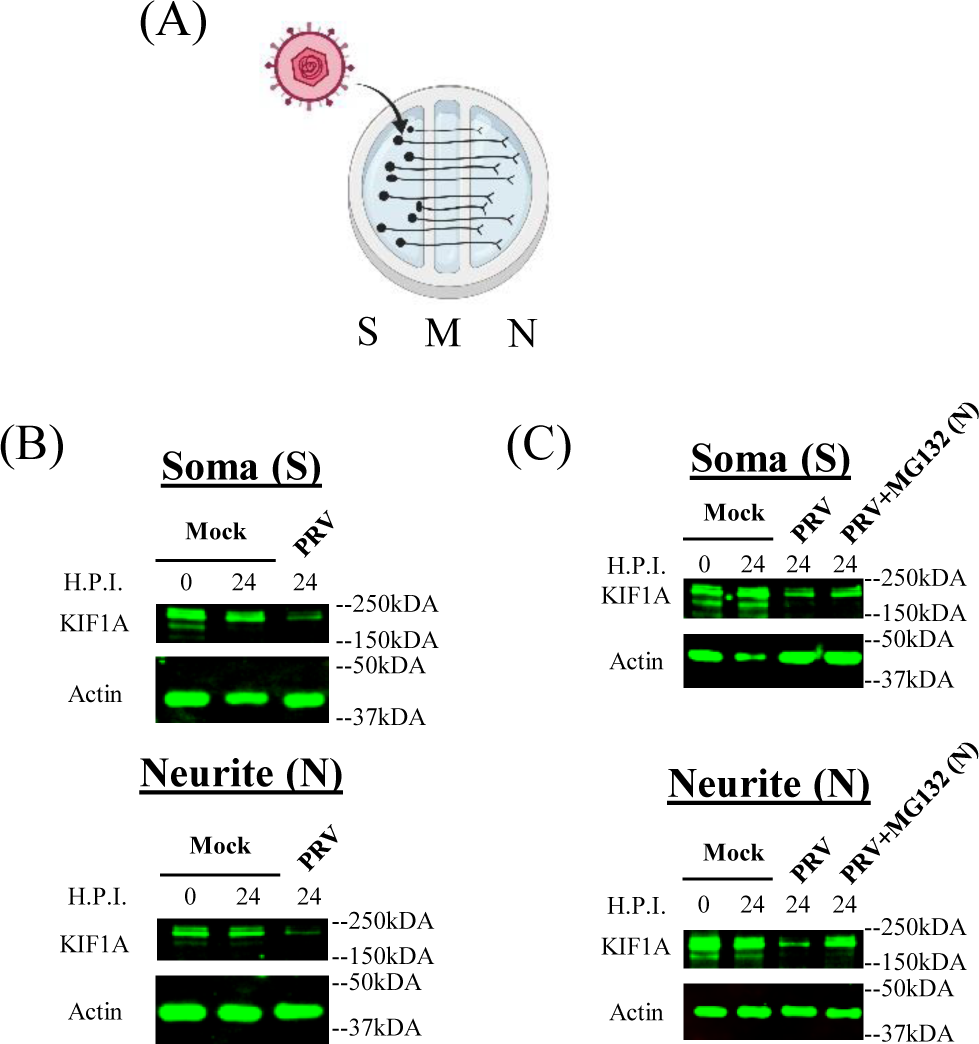
PRV infection of compartmented primary neuronal cultures leads to reduction of KIF1A protein and proteasomal degradation in axons. (A) Illustration of *in vitro* culture of primary superior cervical ganglion neurons using modified Campenot chambers that include the S (soma), M (methocel), and N (neurite, axon termini) compartments. 10^6 PFU of PRV Becker was added in the S (soma) compartment. (B and C) SCG neurons were infected at the Soma (S) compartment with PRV Becker for 24 hours. (B) Cell bodies and axons from S and N compartments were collected at the time intervals show, and lysate were analyzed by Western blot to monitor the levels of KIF1A and Actin. (C) DMSO or MG132 (2.5 μM) were added to the N (Neurite) compartment at 8 hpi. Cell bodies and axons from S and N compartments were collected at the time intervals show, and lysate were analyzed by Western blot to monitor the levels of KIF1A and Actin.

We further determined if the loss of KIF1A protein is mediated by proteasomes in isolated SCG axons during infection. SCG neurons cultured in trichambers were infected with PRV Becker in the S compartment, followed by MG132 treatment in the N compartment at 8 hpi. KIF1A protein levels were reduced significantly in both S and N compartments after PRV Becker infection (Figure 5C). Moreover, axonal KIF1A levels were partially restored by MG132 treatment in the N-compartment (Figure 5C). We concluded that PRV infection accelerates the degradation of KIF1A proteins by proteasomes in axons.

### Expression of both PRV early and late genes are involved in accelerated KIF1A degradation during infection

We next searched for viral proteins that might accelerate KIF1A protein degradation. First, we determined if early events such as virion entry and tegument protein delivery would be enough to promote KIF1A degradation. We infected PC12 cells either with UV treated PRV Becker virions (UVPRV) or with a PRV mutant that lacks IE180 (PRV HKO146), the single master immediate early transcriptional activator of viral genes required for RNA transcription and DNA replication (28, 29). Both UVPRV and PRV HKO146 (grown in IE180-complementing cells) enter the cells, release outer tegument proteins (including vhs, the virion host shutoff protein), and deposit viral DNA in the nucleus, but neither can initiate *de novo* viral protein synthesis. We found that after either infection, KIF1A was not degraded (Figure 6A). We concluded that PRV entry into PC12 cells is not sufficient to accelerate KIF1A protein degradation.

**Fig. 6.**
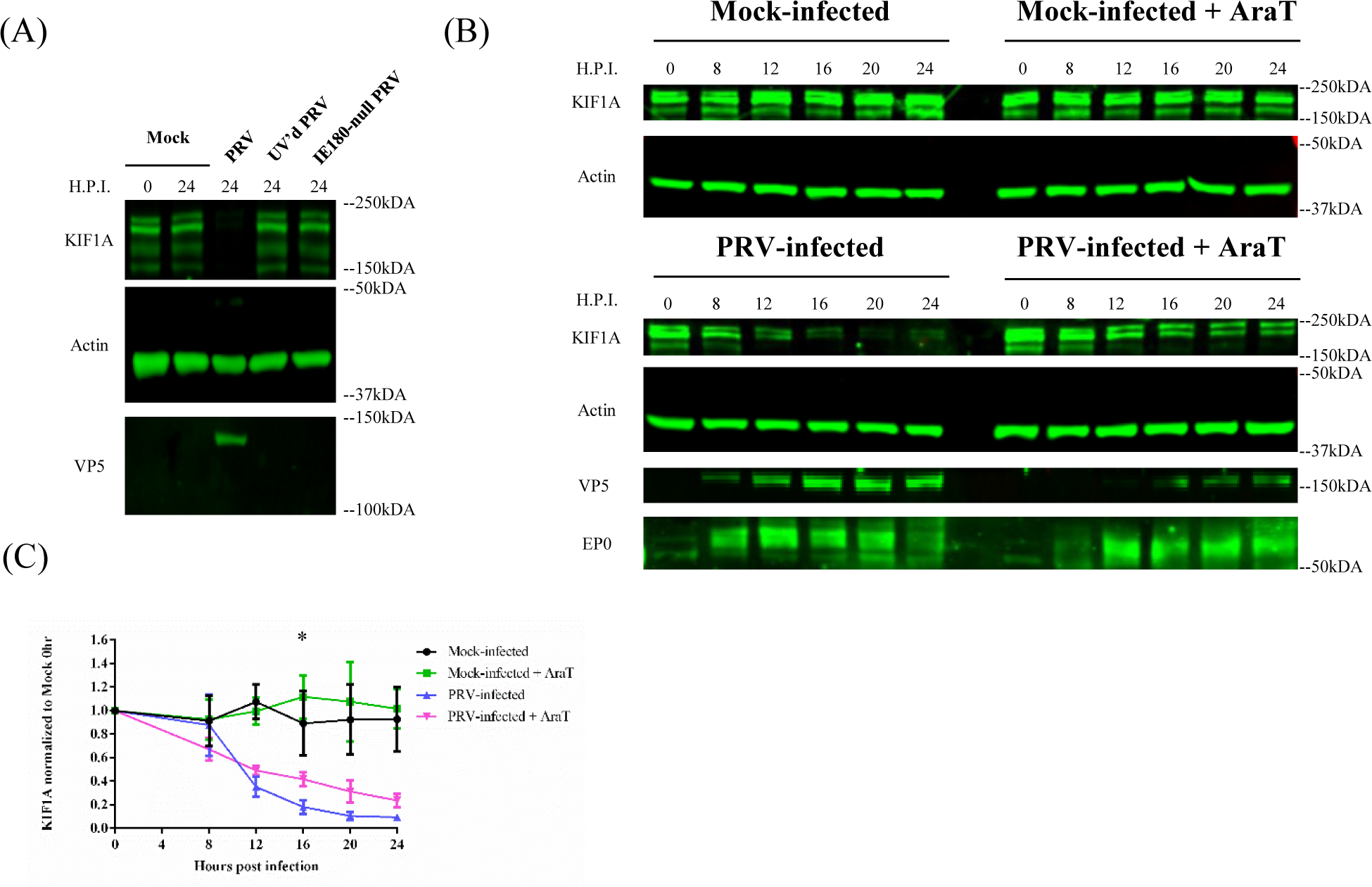
Accelerated proteasomal degradation of KIF1A during infection requires the expression of both PRV early and late proteins. (A) *De novo* PRV viral protein synthesis is required for KIF1A degradation. Differentiated PC12 cells were mock-infected or infected with PRV Becker, UV’d PRV Becker, or IE180-null PRV (HKO146) for 24 hours with MOI of 20. Cells were harvested and lysates were analyzed by Western blot to monitor the levels of KIF1A, actin, and VP5 at indicated time intervals. (B) Differentiated PC12 cells were mock-treated or treated with AraT (100μg/ml) starting 15 minutes before being infected with PRV Becker for 0, 8, 12, 16, 20, 24 hours. Samples were collected at the indicated time intervals. Lysates were subjected to Western Blot analysis to measure protein levels of KIF1A, actin, VP5, and EP0. (C) For every infection, KIF1A protein abundance were measured by band intensities and normalized with respect to actin levels at each time interval. Normalized values were then normalized to that for Mock 0 hour samples. Values are means plus SEMs (error bars) from three independent experiments. *, P<0.05 for PRV Becker vs. PRV Becker with AraT comparison at specified time interval.

We next determined if PRV immediate-early and early, or late genes were involved in KIF1A protein degradation. We pretreated differentiated PC12 cells with the herpesvirus DNA replication inhibitor AraT for 15 minutes, and infected them with PRV Becker for 8, 12, 16, 20, 24 hours. AraT inhibition should have minimal effects on early viral gene expression, but should severely diminish late gene expression. As expected, we detected comparable amounts of the early protein, EP0, in both AraT treated and untreated cells during infection, whereas the accumulation of the late protein VP5 (also major capsid protein, MCP) was severely affected by the AraT treatment (Figure 6B). AraT treatment led to a minor, though statistically significant, stabilization of KIF1A protein in the later stage of infection (Figure 6B-C). This experiment demonstrated that while PRV IE and/or E gene products could promote a reduction in KIF1A protein levels (e.g. potentially through the reduction of KIF1A transcripts), viral proteins accumulating in the late phase of infection are also required for efficient degradation of KIF1A.

### PRV anterograde-sorting complex US9/gE/gI promotes the accelerated degradation of KIF1A proteins

PRV particles in transport vesicles use KIF1A for sorting into axons and subsequent transport. The recruitment of KIF1A to transport vesicles containing enveloped PRV particles at the TGN requires viral membrane protein US9. This interaction is stabilized by the heterodimer of gE/gI. We had noted previously that two ubiquitin ligases, NEDD4 and viral EP0, co-purified with US9 and KIF1A (14). Since KIF1A instability results from proteasomal degradation, we hypothesized that the tripartite complex US9/gE/gI might be involved in accelerating KIF1A degradation. We infected differentiated PC12 cells with PRV Becker and two PRV mutants that were deleted for US9, gE, and gI genes, and are deficient in axonal sorting and anterograde spread: PRV Bartha and PRV BaBe (PRV Becker with the Us region of Bartha). Infection with either mutant lacking US9, gE, and gI partially restored KIF1A proteins during PRV infection (Figure 7A and B). As both of these mutant viruses degrade host transcripts presumably through early proteins, KIF1A levels show a comparable decrease between 8-12 hpi to Becker infection. However, we did not detect the rapid decrease in KIF1A protein levels after 12 h in Bartha and BaBe infections, suggesting that the tripartite complex help accelerate the degradation of KIF1A by proteasomes (Figure 1A and B). We further infected PC12 cells with two PRV US9 mutants deficient in anterograde spread, PRV 161 (US9-null), and PRV 172 (Y49-50A US9, functionally null). In PRV172 infections, all US9, gE, and gI proteins were present, but the dityrosine mutation within the acidic cluster of US9 gene disrupted the formation of the tripartite complex, and made the virus particles incapable of recruiting KIF1A and anterograde spread (14, 30, 31). Both PRV 161 and PRV 172 infections restored KIF1A proteins in magnitudes comparable to infections with PRV Bartha and Babe (Figure 7A and B). These results suggested that the formation of the tripartite complex is necessary for the accelerated loss of KIF1A proteins.

**Fig. 7.**
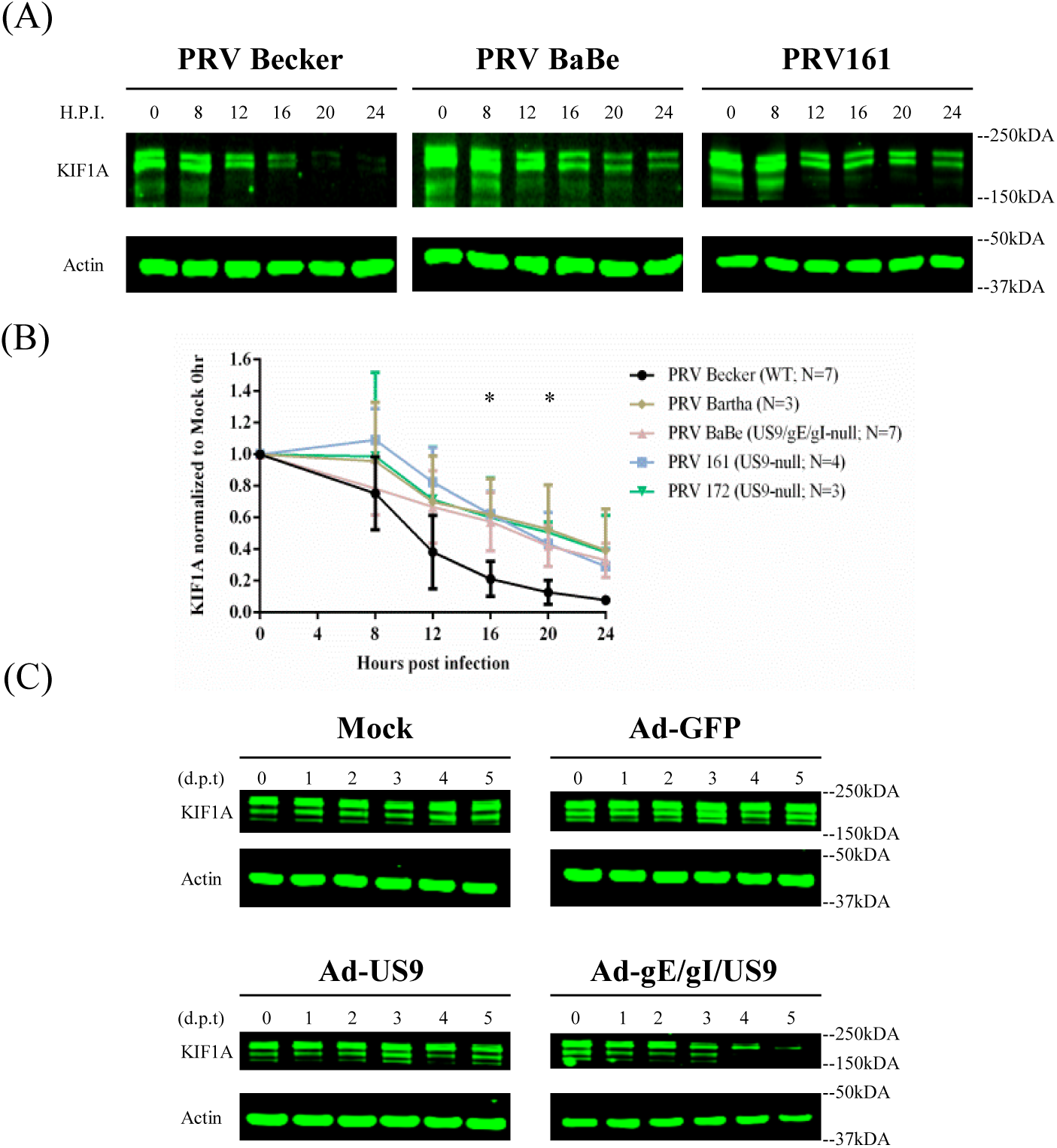
PRV anterograde-spread complex US9/gE/gI promotes the accerlated degradation of KIF1A proteins. (A) Differentiated PC12 cells were infected with PRV Becker, or different PRV mutants defective in anterograde spread (results from PRV BaBe and PRV161 infections are shown) for 0, 8, 12, 16, 20, 24 hours. Cells were harvested and lysates were analyzed by Western blot to monitor KIF1A and actin protein Level. (B) For every infection, KIF1A levels were measured by band intensities and normalized with respect to actin levels at each time interval. Normalized values were then normalized to that for Mock 0 hour samples. Values are means plus SEMs (error bars) from seven independent experiments. *, P<0.05 for all mutants vs. PRV Becker at specified time interval. (C) Differentiated PC12 cells were transduced with Adenoviral vectors expressing GFP, GFP-US9, or GFP-US9/gE-mCherry/gI-mTurquoise 2. Cells were collected at indicated days post transduction (d.p.t.) and subjected to Western Blot Analysis to monitor KIF1A and Actin level.

We further determined if the US9/gE/gI complex could target KIF1A for degradation independent of PRV infection. We transduced differentiated PC12 cells with adenovirus vectors expressing the US9, gE, and gI protein, either individually or in combination, for 1, 2, 3, 4, 5 days. Neither the control Ad-GFP nor Ad-GFP-US9 transduction resulted in KIF1A protein instability, but when PC12 cells were transduced with equal units of the three adenovirus vectors expressing US9, gE, and gI, the KIF1A protein was completely degraded by 5 days post transduction (Figure 7C and Figure S2). We concluded that a functional anterograde sorting complex could augment the proteasomal degradation of KIF1A protein independent of infection by PRV.

Altogether these results suggest that PRV infection leads to the depletion of KIF1A proteins in infected neurons both by blocking new protein synthesis and also specifically targeting the motor for anterograde spread of progeny late in infection, which eventually accelerates proteasomal degradation of KIF1A.

## Discussion

A major distinction of alphaherpesviruses from most of other neuroinvasive viruses is their ability to travel in both anterograde and retrograde directions in axons and in neuronal circuits (4). The anterograde transport of newly assembled herpesvirus virions is crucial for the viral life cycle because it ensures that virus particles are sorted into axons to facilitate inter-host transmission. KIF1A mediates the sorting and transport of PRV egressing virions in axons, with the dynamics of transport involving long segment runs with high velocity (Vmax ∼2.4μm/s) (16). The anterograde transport of PRV virions was greatly hindered in the neurons expressing the dominant-negative form of KIF1A protein (14). In this study, we demonstrated that KIF1A is intrinsically prone to degradation by the proteasomes, and that PRV infection accelerates such degradation and results in near depletion late in infection. The steady state concentration of KIF1A protein in uninfected neuronal cells is achieved through rapid protein synthesis and proteasomal degradation. Upon PRV infection, KIF1A mRNA was degraded in a nonspecific manner, reducing KIF1A protein synthesis while the existing proteins were gradually degraded by the proteasome. In addition, adenovirus vectors expressing proteins constituting the PRV anterograde sorting complex US9/gE/gI induced a decline in KIF1A protein level independent of PRV infection, suggesting the US9/gE/gI tripartite complex accelerates the proteasomal degradation of KIF1A and contributes, alongside PRV host shutoff, to the decline in KIF1A protein during PRV infection. Importantly, using compartmented primary neuron cultures we showed that proteasomal degradation of KIF1A is accelerated in axons after PRV infection.

### The intrinsic instability of the KIF1A protein

The kinesin motor KIF1A mediates fast axonal transport of membranous vesicles including synaptic vesicle precursors and dense core vesicles, and is critical in synaptogenesis (17, 18). In differentiated PC12 cells, KIF1A protein levels decreased rapidly upon treatment with the protein synthesis inhibitor cycloheximide (CHX), and showed an apparent half-life of approximately 4 hours. The half-lives of kinesin-1 and −2 motors were longer under the same condition (>8 hours). Our results are consistent with a previous proteomics study using rat cortical neurons, where KIF1A had the 8^th^ shortest half-life in among ∼2800 proteins measured (t_1/2_ = 0.42 day) (32), suggesting that this is a consistent characteristic of KIF1A in both CNS and PNS neurons. The decline in KIF1A concentration is blocked when PC12 cells were treated simultaneously with CHX and MG132, demonstrating that the loss of KIF1A protein was predominately mediated by proteasomes. Our results confirmed another previous proteomics study where the half-life of the KIF1A protein increased and the degradation rate dropped upon inhibition of proteasomes (33).

UNC-104, the KIF1A homolog in *C. elegans*, was shown to be degraded at synapse regions *in vivo* in an ubiquitin-dependent manner. Moreover, the stability of the motor was linked to its capacity for cargo binding (22). It was also proposed that the degradation of Liprinα1, an cargo adaptor for KIF1A, by either proteasome or CaMKII phosphorylation was required for the “unloading” of LAR (leukocyte common antigen-related) family of receptor protein tyrosine phosphatases at synapses (34, 35). These studies imply a potential mechanistic link between motor degradation and cargo release. Since we also observed proteasomal degradation of KIF1A in fluidically and physically separated axons during PRV infection, KIF1A may be actively degraded in axons by proteasomes upon release of viral cargo.

The relative stabilities of KIF5 and KIF3A proteins were unexpected, since several lines of evidence suggested that both motors are actively degraded in axons (36–39). However, unlike KIF1A, both KIF5 and KIF3A also mediate short distance transport of cellular cargoes within cell bodies, and may be recycled for future use. The particular short half-life of KIF1A is consistent with the idea that KIF1A is a specialized motor for axonal transport.

### The reduction of KIF1A mRNA upon PRV infection

The dynamic steady state level of KIF1A protein in uninfected cells suggests that, in PRV-infected cells, KIF1A protein concentration may be sensitive to viral host shutoff or mRNA instability induced by infection. Alphaherpesviruses reduce/block the synthesis of some cellular proteins after infection to facilitate viral gene expression and to evade host antiviral responses. The virion host shutoff protein (vhs) is an endoribonuclease encoded by the viral UL41 gene (40–42). HSV-1 vhs enters the infected cells as a part of incoming virion tegument, and subsequently interacts with translation initiation factors eIF4H, eIF4B, and eIF4A to degrade host mRNAs and halts host protein translation (43, 44). Infection of PC12 cells with inactivated virions (UV’d, IE180-null PRV) did not lead to the decline in KIF1A protein. We conclude that tegument proteins like vhs delivered by entering particles are not sufficient for the reduction in KIF1A protein during infection.

We further analyzed the accumulation of cellular mRNAs during infection and compared the levels of kinesin motor transcripts. qRT-PCR analysis of PRV-infected PC12 cells revealed that the levels of KIF1A, KIF5B, and GAPDH transcripts all decreased significantly at similar rates, starting as early as 3 h.p.i., and reaching almost full depletion at 12 h.p.i. The reduction of KIF1A mRNA reduced the rate of new protein synthesis. This reduction of new KIF1A proteins, coupled with the active degradation of existing KIF1A motors by proteasomes, leads to the significant decrease in KIF1A protein concentration in infected neuronal cells.

### The PRV anterograde sorting complex US9/gE/gI directs more KIF1A motors to be degraded by proteasomes

US9 forms a complex with two other membrane proteins, gE and gI to sort progeny PRV virions in transport vesicles into axons (14, 15)(J. Scherer, I. B. Hogue, Z. A. Yaffe, N. S. Tanneti, B. Y. Winer, M. Vershinin, L. W. Enquist, submitted for publication). We have previously identified KIF1A as an interaction partner of PRV US9 protein (14). In addition, we had noted that two ubiquitin ligases, NEDD4 and viral EP0, co-purified with US9 and KIF1A (14). Since we also showed accelerated proteasomal degradation of KIF1A during infection, we hypothesized that the PRV anterograde sorting complex, gE/gI/US9, could independently recruit KIF1A to viral transport vesicles for axonal sorting and transport, and eventually promote the proteasomal degradation of KIF1A in axons. We investigated this hypothesis using adenovirus vectors to express the components of the anterograde sorting complex (one by one and in combination) in neuronal cells without PRV infection. In cells transduced with adenovirus vectors expressing US9, gE, and gI, US9 interacts with the gE/gI heterodimer in lipid rafts, and the US9, gE, and gI complex was observed to undergo axonal anterograde transport together with long-range motility characteristic of anterogradely-moving PRV virions in axons (J. Scherer, I. B. Hogue, Z. A. Yaffe, N. S. Tanneti, B. Y. Winer, M. Vershinin, L. W. Enquist, submitted for publication). PRV envelope proteins, US9/gE/gI, when expressed together in neuronal cells, induced a significant decline in KIF1A protein concentration. Therefore we concluded that the viral anterograde sorting complex US9/gE/gI promotes the accelerated loss of KIF1A proteins.

This result is consistent with experiments where PC12 cells were infected with PRV strains that fail to express a functional US9/gE/gI complex to recruit KIF1A to undergo anterograde spread. These PRV mutants still promote host shutoff, leading to the decline of KIF1A proteins late in infection. However, the rate of decline is slower than that observed after wildtype PRV infection, suggesting that the US9/gE/gI complex accelerates the degradation of KIF1A by the proteasomes. The inhibition of late gene expression (e.g. US9) by AraT treatment also led to a minor yet statistically significant stabilization of KIF1A protein concentration. We concluded that the PRV anterograde sorting complex US9/gE/gI directs more KIF1A motors to be degraded by proteasomes. This might shed light on the mechanisms underlying the intrinsic instability of KIF1A protein, which potentially stem from the interaction of the motor with its cellular cargoes.

### PRV infection induces the accelerated loss of KIF1A protein through two separate mechanisms

Our model to account for our results is shown in Figure 8. KIF1A motors, mediating the anterograde transport of the vesicles in axons critical for synapse formation and function, are inherently prone to proteasomal degradation. Such intrinsic instability results in the dynamic steady state level of KIF1A protein maintained through rapid protein synthesis and proteasomal degradation in neuronal cells (Fig. 8A). The balance between protein synthesis and degradation is disrupted by PRV infection through two separate mechanisms (Fig. 8B). First, PRV infection promotes the nonspecific loss of KIF1A mRNAs, and reduces the synthesis of KIF1A protein. The concentration of existing KIF1A protein continues to decline through default proteasomal degradation. Second, PRV infection accelerates the degradation of KIF1A protein through the PRV anterograde sorting complex US9/gE/gI. We propose that US9 recruits KIF1A to transport vesicles containing enveloped virions or light particles, as well as vesicles with viral glycoproteins for axonal sorting and transport, and eventually this complex promotes the proteasomal degradation of KIF1A in axons.

**Fig. 8.**
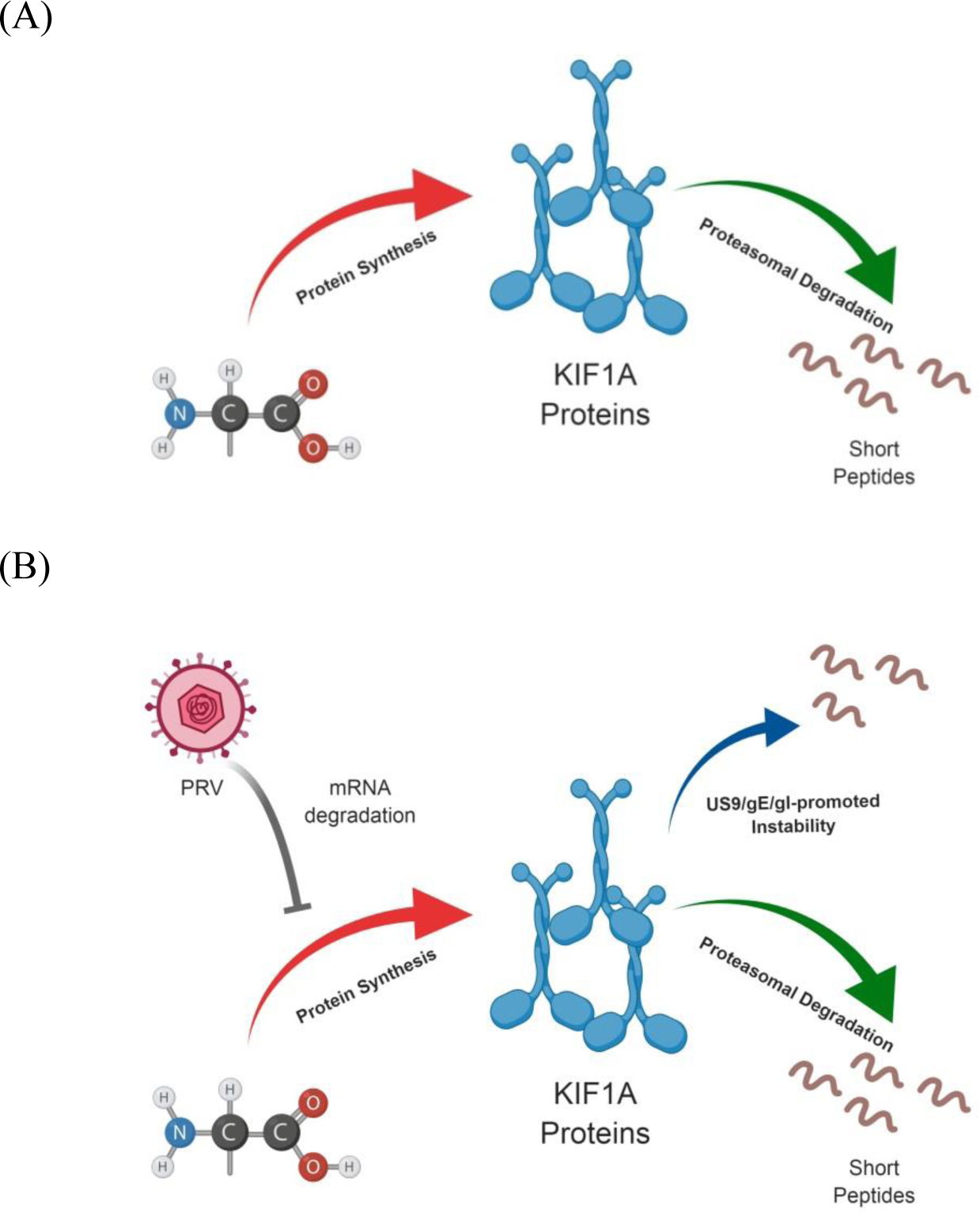
PRV infection induces the accelerated loss of KIF1A protein through two separate mechanisms. (A) In uninfected cells, KIF1A protein is inherently prone to be degraded by the proteasome. The steady state concentration of KIF1A protein is achieved through rapid protein synthesis and proteasomal degradation. (B) PRV infection induces KIF1A mRNA degradation and reduced protein synthesis, while the existing KIF1A proteins undergo default proteasomal degradation. Furthermore, the proteasomal degradation of the existing KIF1A proteins is accelerated by the PRV gE/gI/US9 complex in the late phase of infection. Images were created with BioRender.

Our work implies that after reactivation from latently infected PNS neurons, the targeted loss of KIF1A motor proteins leads to a limited time window for sorting into and transport of viral particles in axons for further host-to-host spread. Indeed, the loss of available KIF1A motors over the course of PRV infection results in reduced recruitment of the motor by US9 (14). While our work has focused on PRV, it would be important to know if infection by other alphaherpesviruses promotes differential degradation of axonal motor proteins. The anterograde transport of HSV-1 has been shown to be mediated by kinesin-1 (KIF5A, KIF5B, and KIF5C) and kinesin-3 (KIF1A) motors (11, 45)(E. Engel and L.W. Enquist, unpublished data). It is therefore of clinical importance to determine if HSV-1 infection induces similar loss in human neurons *in vitro*. Such a loss of kinesin-1 and kinesin-3 motors after reactivation from PNS neurson could have significant consequence for long term neuronal function. Further work is required to understand whether PNS neurons can tolerate the loss of specific kinesin motors during reactivation episodes, and whether the motor concentrations are restored after reactivation if those neurons survive.

## Materials and Methods

### Cell lines and viruses

Porcine kidney epithelial cells (PK15, ATCC) were maintained in Dulbecco modified Eagle medium (DMEM, Hyclone) supplemented with 10% fetal bovine serum (FBS, Hyclone) and 1% penicillin-streptomycin (Hyclone). PK15 cells were used to propagate and determine titers of all PRV strains used in this study. 293A cells (Invitrogen) were used to propagate and determine the titer of adenovirus vectors used in this study. The neuronal PC12 cell line has been used extensively to model primary neurons infection of PRV and to reproduce the US9-associated transport phenotype (14, 46). PC12 cells were cultured on dishes coated with type 1 rat tail collagen (BD Bioscience, Bedford, MA) in 85% RPMI 1640 (Thermo Fisher) supplemented with 10% horse serum(Life Technologies) and 5% FBS (47). PC12 cells were differentiated in RPMI 1640 supplemented with 1% horse serum and nerve growth factor (NGF, 2,5S, Invitrogen) at 100 ng/ml (47). Differentiation medium was replaced every third day for eight days before infection.

The PRV viral strains used in this study include both wild-type laboratory stain and recombinants. PRV Becker is a wildtype strain (25, 26). PRV Bartha is an attenuated vaccine strain (48, 49). PRV BaBe is a viral recombinant in the PRV Becker background with deletions from the unique short (US) region of PRV Bartha (50). HKO146 lacks the only PRV immediate early gene IE180 and expresses hSyn Cre-t2a-venus in US9 loci in the PRV Becker background (H. Oyibo, P. Znamenskiy, H. V. Oviedo, L. W. Enquist, A. Zador, unpublished data). PRV161 harbors a complete deletion of the US9 gene in the PRV Becker background (51). PRV172 harbors two tyrosine to alanine substitutions (Y49 Y50 to AA) in the PRV Becker background (30). Individual replication-deficient adenovirus vectors expressing GFP, GFP-US9, gE-mCherry, gI-mTurquoise 2 were previously reported (14, 52).

### Primary neuronal culture

Embryonic superior cervical ganglia (SCG) neurons were isolated and cultured as previously described (53). Briefly, superior cervical ganglia (SCG) were isolated from day 17 Sprague-Dawley rat (Hilltop Labs) embryos, plated and maintained on poly-DL-ornithine (Sigma-Aldrich) and laminin (Invitrogen) coated dishes (Falcon or MatTek) in neuronal medium made with neurobasal medium (Gibco) supplemented with 1% penicillin-streptomycin with 2 mM glutamine (Invitrogen), 2% B-27 (Gibco), and 100 ng/ml neuronal growth factor (NGF, 2.5S (Invitrogen)). On two days post plating, 1 mM cytosine-D-arabinofuranoside (AraC; Sigma-Aldrich) was added to selectively eliminate dividing non-neuronal cells in culture for two days. SCG neurons were allowed to differentiate for at least 14 days before infection.

Compartmented neuronal cultures were prepared on poly-ornithine and laminin coated plastic dishes. Parallel grooves were etched into the plastic dish, and 1% methylcellulose in neuronal medium was added to the region that would underlie the middle compartment of the isolator rings. CAMP320 isolator rings (Tyler Research) were coated with autoclaved silicone vacuum grease on one side and gently applied to the dish so that the etched grooves extended across three compartments. Dissociated SCG neurons were plated in the soma compartment (S), and were allowed to extend axons along the grooves through the M compartment and into the neurite (N) compartment for 14-28 days.

### Viral infection in neuronal culture

Triplicates of differentiated PC12 cells were infected with the indicated PRV strain at a multiplicity of infection (MOI) of 20 at the indicated hours post infection. Infected cells from each triplicate were harvested and combined into a single sample for further biochemical analysis. Compartmented SCG neuronal cultures were infected in the soma (S) compartment for 24 hours. Neuronal cell bodies from the S compartments and axons from the N compartments were harvested for biochemical analysis. The proteasome inhibitor MG132 (Millipore Sigma) was prepared in DMSO and added at a concentration of 2.5 µM. The herpesvirus DNA replication inhibitor thymine 1-β-D-arabinofuranoside (AraT) (Millipore Sigma) was added at a concentration of 100μg/ml 15 minutes prior to infection.

### Adenovirus vectors transduction

The adenovirus vectors expressing GFP, GFP-US9, gE- mCherry, and gI-mTurquoise-2 were reported previously (14, 52). Adenovirus vectors were propagated in 293A cells, and cell-associated virus was harvested in serum-free DMEM media. Adenoviral transduction of differentiated PC12 cells was performed for up to five days to ensure the expression of fluorescent transgene(s) in the majority of cells (>90%).

### Quantitative Western blotting analysis

Neuronal cell lysates were prepared in radioimmunoprecipitation assay buffer (RIPA) (50mM Tris-HCl, 150mM NaCl, 5mM EDTA, 1% NP-40, 0.1% SDS, 0.1% Triton X-100, and 1% sodium deoxycholate, pH 8.0) supplemented with 1 mM dithiothreitol (DTT, Sigma-Aldrich) and protease inhibitor cocktail (Sigma-Aldrich). Cell lysates were kept on ice for one hour and centrifuged at 13,200 rpm at 4°C for 10 minutes. Cleared cell lysates were transferred to a new tube and mixed with 4X LDS sample loading buffer (Invitrogen) supplemented with 240 mM DTT. The samples were heated at 95°C for 10 minutes and centrifuged at 4°C for 10 minutes to remove potential protein aggregates before proteins were separated in gradient polyacrylamide gels (4-12%) (Invitrogen). Proteins were transferred onto nitrocellulose membranes (Whatman) by semidry transfer (Bio-Rad). Membranes were blocked in 5% nonfat dry milk in Tris buffer saline with 0.1% Tween 20 (TBS-T) for 1 hour at room temperature. Primary and secondary antibodies for immunoblot analysis were prepared in 1% milk in TBS-T. Protein bands were imaged by LI-COR Odyssey CLx imaging system. The signal intensity was quantified using LI-COR Image Studio Lite software.

Antibodies used in this study included a mouse monoclonal KIF1A antibody (Clone 16, BD, at 1:2000), a mouse monoclonal kinesin heavy chain antibody (MAB-1614, Millipore Sigma, at 1:1000), a mouse monoclonal KIF3A antibody (Clone K2.4, Covance Inc., at 1:1000), a mouse monoclonal β-Actin antibody (Sigma, at 1:10000), a monoclonal US9 antibody (IA8, DSHB, at 1:100), a rabbit polyclonal GST-EP0 antibody at 1:2000 (54), a mouse monoclonal PRV VP5 antibody at 1:1000 (55), a mouse monoclonal PRV US3 antibody at 1:10000 (56), a rabbit polyclonal gE cytoplasmic tail antibody at 1:1000 (57), a rabbit polyclonal gI antibody, a generous gift from K. Bienkowska-Szewczyk, at 1:1000 (58). IRDye secondary antibodies included 800CW donkey anti-mouse IgG, 800CW donkey anti-rabbit IgG, and 680RD donkey anti-rabbit at 1:15000 (LI-COR).

### qRT-PCR analysis

Differentiated PC12 cells infected with PRV were harvested and subjected to total RNA extraction using an RNeasy Plus Mini Kit (Qiagen). First strand cDNA synthesis of isolated RNAs was performed with SuperScript III First Strand synthesis kit with Oligo (dT) primer (Invitrogen). Quantitative reverse transcription PCR was performed using Eppendorf Realplex^2^ Mastercycler with reaction mixtures prepared with KAPA SYBR Fast Universal qPCR Kit (KAPA Biosystems). The PCR primers for KIF1A, KIF5B, GAPDH, and 18s rRNA were reported elsewhere (59). The qRT-PCR reaction samples were prepared in triplicates. The relative abundance of RNA in each sample was calculated using the –ΔΔC_T_ method normalized to 18s rRNA and subsequently to uninfected samples.

### Statistical analysis

One-way analysis of variance (ANOVA) with Tukey’s posttest, and two-way ANOVA with Sidak’s posttest were performed using GraphPad Prism 6. Values in text, graphs, and figure legends throughout the manuscript are means ± standard errors of the means (SEM).

### Image production

Cartoon illustrations were made with web application BioRender under Princeton University license for education use.

### Ethics Statement

All animal work was performed in accordance with the Princeton Institutional Animal Care and Use Committee (protocols 1947-16). Princeton personnel are required to adhere to applicable federal, state, local and institutional laws and policies governing animal research, including the Animal Welfare Act and Regulations (AWA); the Public Health Service Policy on Humane Care and Use of Laboratory Animals; the Principles for the Utilization and Care of Vertebrate Animals Used in Testing, Research and Training; and the Health Research Extension Act of 1985.

## Acknowledgement

We thank Halina Staniszewska-Goraczniak for excellent technical and logistical support, and the rest members of the Enquist Lab for their critical comments on the projects. This project was funded by NIH, National Institute of Neurological Disorders and Stroke (NINDS) RO1 NS033506 and RO1 NS060699 (L.W.E.).

---

**Figure S1.**
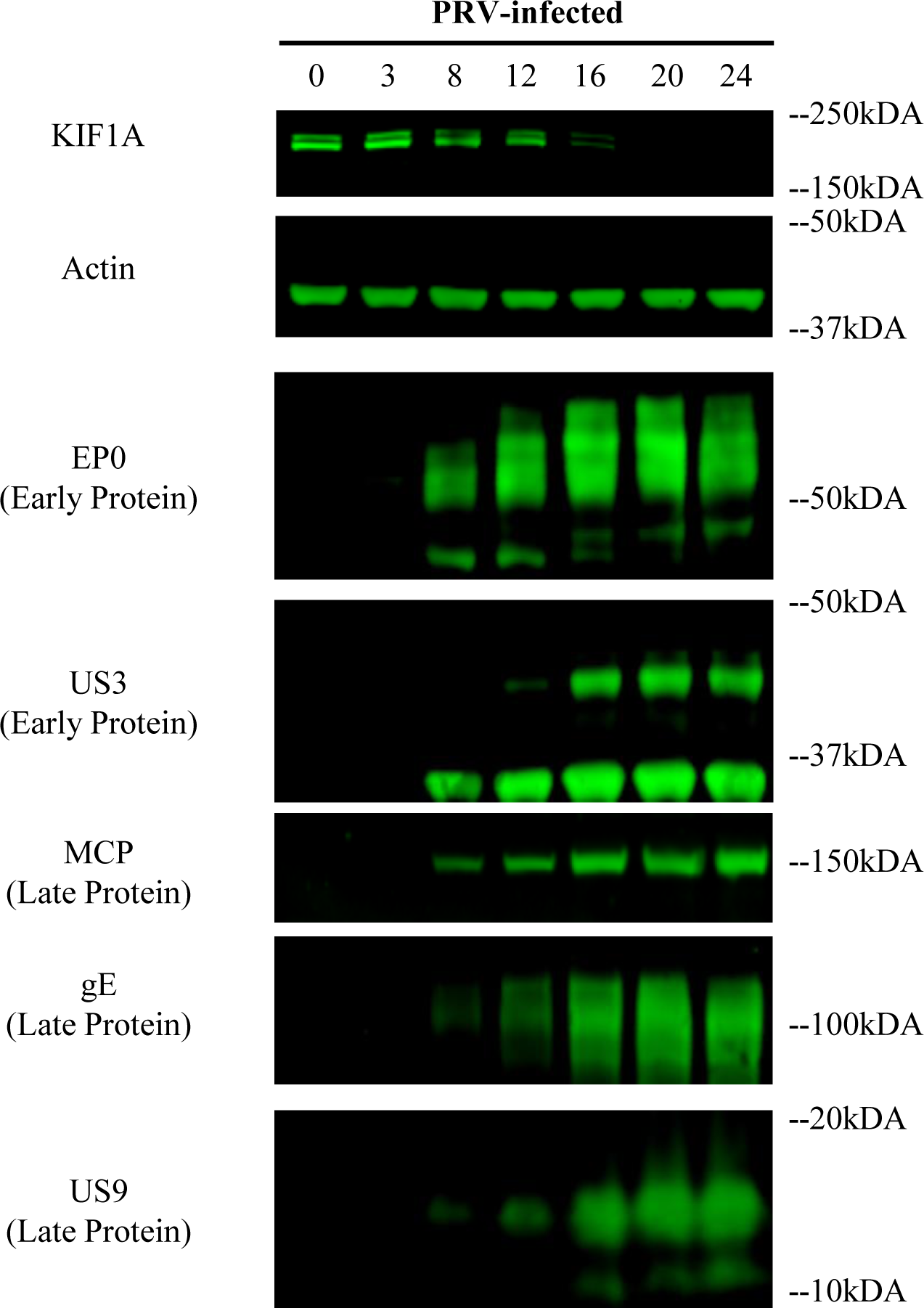
PRV early and late gene expressions in differentiated PC12 cells.

**Figure S2.**
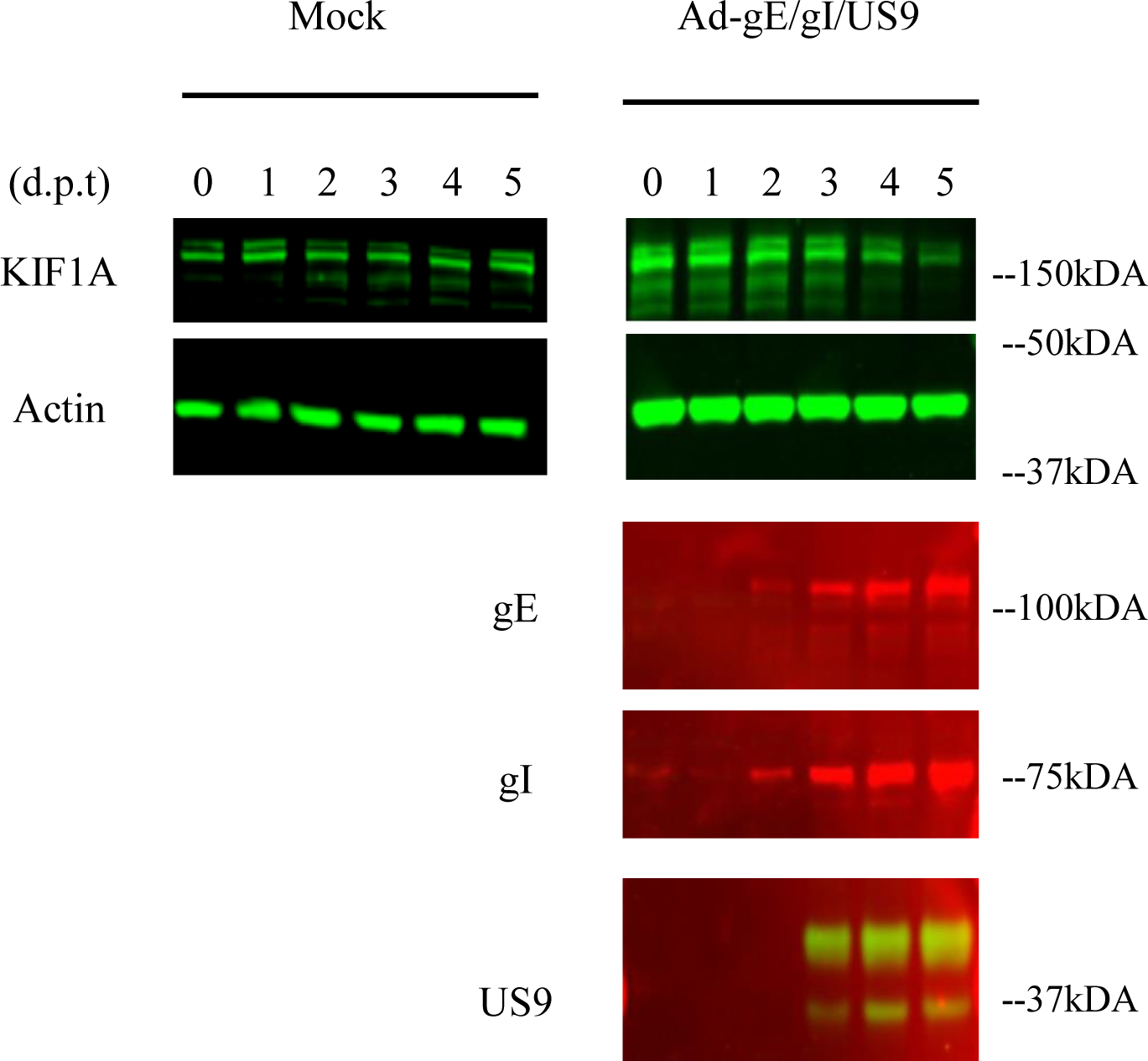
**Expression of transduced genes US9, gE, and differentiated PC12 cells.**

